# Development and pilot screen of novel high content assay for down regulators of expression of heterogenous nuclear ribonuclear protein H2

**DOI:** 10.1101/2020.10.05.326116

**Authors:** Juan Diez, Sumitha Rajendrarao, Shadi A. Baajour, Praathibha Sripadhan, Timothy P. Spicer, Louis D. Scampavia, Dmitriy Minond

## Abstract

Despite recent advances in melanoma drug discovery, the average overall survival of patients with late stage metastatic melanoma is approximately 3 years, suggesting a need for new approaches and melanoma therapeutic targets. Previously we identified heterogeneous nuclear ribonucleoprotein H2 as a potential target of anti-melanoma compound 2155-14 (Palrasu *et al*, *Cell Physiol Biochem* 2019;53:656-86). In the present study, we endeavored to develop an assay to enable a high throughput screening campaign to identify drug-like molecules acting via down regulation of heterogeneous nuclear ribonucleoprotein H that can be used for melanoma therapy and research.

**Results:** We established a cell-based platform using metastatic melanoma cell line WM266-4 expressing hnRNPH2 conjugated with green fluorescent protein to enable assay development and screening. High Content Screening assay was developed and validated in 384 well plate format, followed by miniaturization to 1,536 well plate format. All plate-based QC parameters were acceptable: %CV = 6.7±0.3, S/B = 21±2.1, Z’ = 0.75±0.04. Pilot screen of FDA-approved drug library (n=1,400 compounds) demonstrated hit rate of 0.5%. Two compounds demonstrated pharmacological response and were authenticated by western blot analysis.

**Conclusions:** We developed a highly robust HTS-amenable high content screening assay capable of monitoring down regulation of hnRNPH2. This assay is thus capable of identifying authentic down regulators of hnRNPH1 and 2 in a large compound collection and, therefore, is amenable to a large-scale screening effort.

## INTRODUCTION

There is an unmet need for the discovery of new molecules that can be targeted to induce melanoma cell-specific death. Several decades of research into kinase inhibitors and immunotherapy have led to the development of a number of new drugs along with the resulting extension of patient lives, but resistance often occurs and it appears that multiple mechanisms need to be targeted in order to provide sustained disease control (1, 2).

Our recent report suggests that spliceosomal protein hnRNPH2 can be targeted to induce melanoma-specific cell death that can address an unmet need for such novel therapeutic approaches focused on novel mechanisms (3).

Targeting multiple pathways may help prevent or overcome drug resistance (4–6). Modulation of hnRNPH2 induces an autophagy which was shown to be an effective strategy in overcoming multi-drug resistance (MDR) in cancers (7). Autophagy plays a role in the development as well as overcoming drug resistance in melanoma (8), therefore, an ability to shift balance from pro- to anti-resistance can also be beneficial. Because of this ability, hnRNPH2 modulation may also address the lack of therapies for NRAS mutants and other subtypes of melanoma. Autophagy also regulates cytokine secretion and antigen presentation and, therefore, contributes to anti-tumor immune responses (9–11).

Based on these considerations, modulation of hnRNPH2 may overcome drug resistant melanoma and help prevent occurrence of resistance, which would be a significant improvement over existing drugs such as vemurafenib.

With this in mind, we decided to explore existing chemical diversity in order to identify novel down regulators of hnRNPH2. Typically, this requires an assay suitable to compound screening in a high throughput format (HTS). However, we were not able to identify an assay platform that would allow us to do so, therefore, we decided to develop such a platform. Herein we present the results of HTS-amenable assay development and its validation.

## MATERIALS AND METHODS

### WM266-4-H2GFP cell line establishment

WM266-4 cells (12) were purchased from Rockland Immunochemicals, Inc. (Limerick, PA) and authenticated by the vendor. Subsequent authentications and mycoplasma certification was done at Johns Hopkins University Genetic Resources Core Facility. WM266-4 cells were plated in 6-well plate at 5 × 10^5^ cells/well. After cells were attached (next day) they were transfected with GFP-H2 plasmid (Sino Biologics) using Lipofectamine Plus according to the manufacturer directions. Briefly, 2 μL of plasmid, 5 μ L lipofectamine and 2.5 μL Plus reagent were mixed in 500 μL serum-free EMEM and after 5 min incubation at RT 250 μL were added to each well. Transfection efficiency was assessed by monitoring green fluorescence. When cells were confluent, they were transferred to a T75 flask and after they adhered selection began by adding 200 μg/mL Hygromycin to the medium. As cells began dying, the medium was frequently changed, maintaining the hygromycin concentration. Two weeks later, when colonies were observed, they were transferred to 6-well plate wells, allowed to grow and then diluted and plated at 1 cell/well in 48-well plates.

Hygromycin was always maintained to keep the selection pressure. The colony with the strongest and most uniform expression was selected and H2-GFP expression was assessed by fluorescent microscopy and Western blot. Cells from this colony were sorted, using FACS to further purify the colony.

### Primary WM266-4-H2GFP HCS assay in 384 well plate format

1,250 WM266-4-H2GFP cells were plated in low volume black sterile tissue culture treated 384-well plates (Greiner cat# 781099) in 10 μL of media. Next day after plating, compounds from FDA approved drug library and pharmacological control (dronedarone) were added to the cells (100 nL per well, 10μM final assay concentration) using the 384 channel pintool (V&P Scientific, San Diego) mounted on ViaFlo 384 (Integra). After overnight incubation at 37°C, 5% CO_2_ and 95% relative humidity, the plates were imaged using algorithm on CellInsight CX7 (Thermo Scientific). For each well, the Average Total Object Intensity (ATOI) was measured and used as a basis for calculation of statistic parameters (whole plate average ATOI, %CV, Z’) and EC_50_ values.

### Primary WM266-4-H2GFP HCS assay in 1,536 well plate format

750 WM266-4-H2GFP cells were plated in low volume black, clear bottom sterile tissue low base culture treated 1,536-well plates (Aurora cat# 00019326) in 5 μL of media using a Flying Reagent Dispenser (FRD, Aurora Biosciences).

The next day, cells were pinned with 50nL of the pharmacological control compounds mitomycin and dronedarone using a pintool transfer unit (Kalypsys/GNF). After overnight incubation at 37°C, 5% CO_2_ and 95% relative humidity, fluorescent images were acquired on CellInsight image reader using 10x objective collecting 4 fields of view per well (485 nm at a fixed exposure time 0.063 s). We implemented the CellInsight’s target activation assay algorithm which was configured for the screen. Background from the raw images was removed by using low pass filter and smoothing parameters. Primary objects were identified using the fixed threshold method set at 65 and the objects touching the image border were excluded. Further, primary objects were validated by defining the object area and intensity of the objects. The output features such as valid object count and the average total intensity were exported for each plate and data analysis was performed using the ratio of valid object count to Average Total Object Intensity.

### GFP interference assay

This assay was conducted the same way as a primary HCS assay except forcompound incubation time. Briefly, WM266-4-H2GFP cells were plated in 384-well plates (Greiner cat# 781099) in 10 μL of media. Next day after plating, test compounds were added to the cells (100 nL per well) using the 384 channel pintool (V&P Scientific, San Diego) mounted on ViaFlo 384 (Integra) and fluorescence emission was scanned immediately using microplate reader Synergy H1 (Biotek).

### Cell viability assays

Briefly, WM-266-4 and WM266-4-H2GFP cells were plated in 384-well plates in 8 μL of media. Test compounds and dabrafenib (pharmacological assay control) were prepared as 10-point, 1:3 serial dilutions starting at 300 μM, then added to the cells (4 μL per well) using the 384 channel pintool (V&P Scientific, San Diego) mounted on ViaFlo 384 (Integra). Plates were incubated for 72 h at 37 °C, 5% CO_2_ and 95% relative humidity. After incubation, 4 μL of CellTiter-Glo® (Promega cat#: G7570) were added to each well and incubated for 15 min at room temperature. Luminescence was recorded using a Biotek Synergy H1 multimode microplate reader. Viability was expressed as a percentage relative to wells containing media only (0%) and wells containing cells treated with DMSO only (100%). Three parameters were calculated on a per-plate basis: (a) the signal-to-background ratio (S/B); (b) the coefficient for variation [CV; CV = (standard deviation/mean) x 100)] for all compound test wells; and (c) the Z’-factor (18).

### Western blotting for hnRNP H2

WM-266-4 and WM266-4-H2GFP cells were sonicated in in RIPA lysis buffer containing protease and phosphatase inhibitors. Protein isolates were subjected to SDS-PAGE followed by transfer to nitrocellulose membrane. hnRNP H2 was detected using rabbit polyclonal antibody (Abgent #: AP13497b; 1:3,000, in 5% milk overnight). After washing with TBST, the membranes were treated with chemiluminescent horseradish peroxidase detection reagent (Thermo Scientific, Cat# 32209) and developed using the Licor imaging system. Licor software was used to quantitate the intensity of protein bands. The protein bands were normalized against loading control (β-actin).

## RESULTS

### WM266-4-H2GFP cell line establishment

We used a construct encoding for hnRNPH2 tagged with GFPSpark (Sino Biological HG16541-ANG) to stably transfect WM266-4 cells (Fig. 1A). hnRNPH2-GFP was transfected into WM266-4 cells using Lipofectamine LTX (Life Technologies 15338030) according to the manufacturer directions. Transfected WM266-4 cells were selected by culturing them in EMEM supplemented with hygromycin until most of the cells died and some colonies appeared. Those colonies were allowed to grow, collected and cells diluted and plated again in 96-well plates at a dilution of 1 cell per well to obtain clonal populations of the H2-GFP cells (Fig. 1B). After growing these colonies for several weeks, the expression was monitored by fluorescent microscopy and those colonies with the stronger and uniform expression of H2-GFP were selected (Fig. 1C). To further enrich cell population expressing hnRNPH2-GFP, cells were sorted using FACS and WM266-4 cells as non-GFP-expressing control. Resulting WM266-4-hnRNPH2-GFP (H2-GFP) cells showed morphology, viability and doubling rate similar to parental cells (Fig. 1D). Similarly, to WM266-4 cells H2-GFP cells recovered well after LN_2_ storage. Western blot analysis confirmed expression of H2-GFP (Fig. 1E).

**Figure 1.**
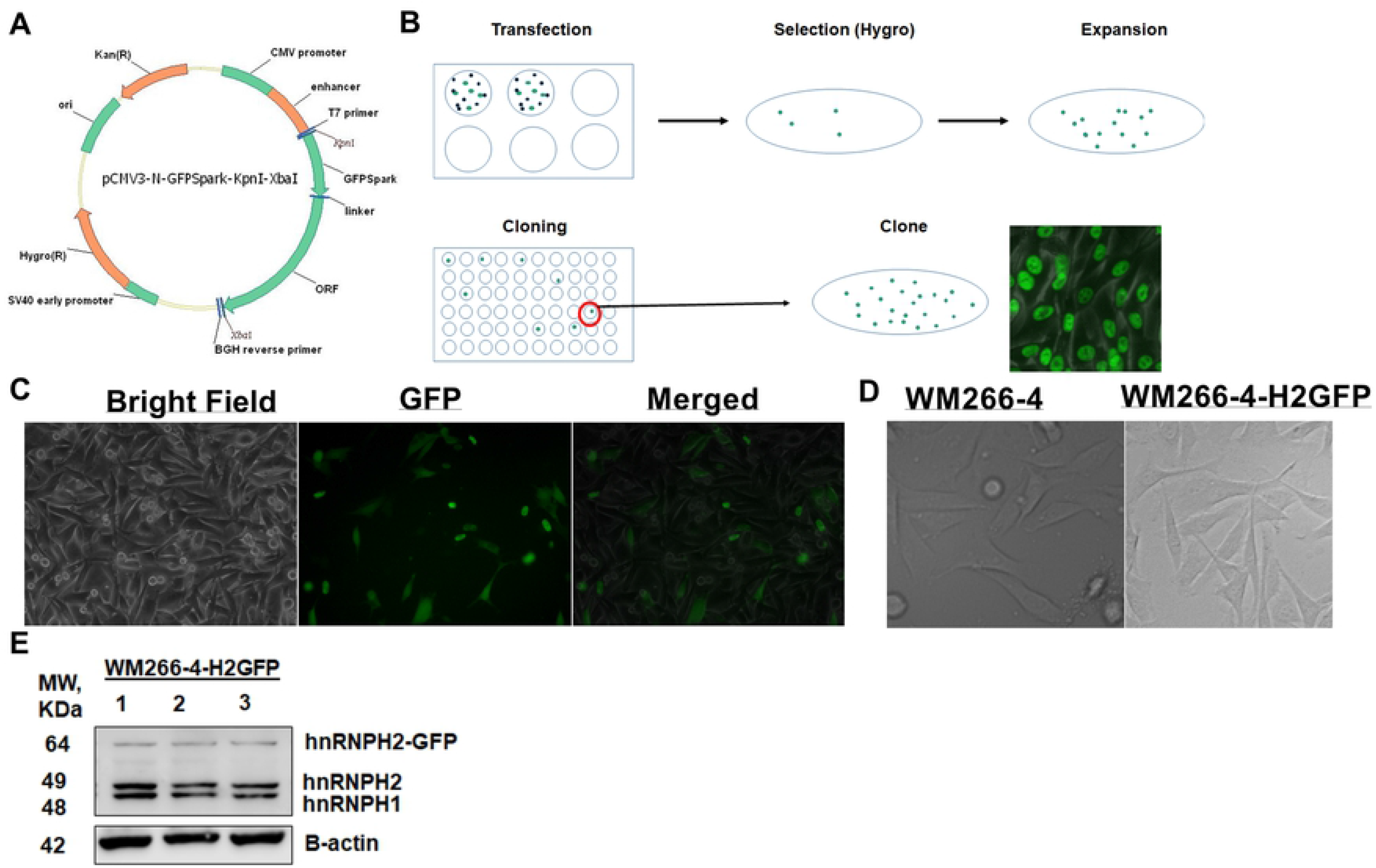
Development of WM266-4-H2GFP cells. (A) Plasmid scheme; (B) WM266-4-H2GFP cell line establishment work flow; (C) Imaging of WM266-4 cells transfected with hnRNPH2-GFP plasmid after clonal expansion; (D) WM266-4 and WM266-4-H2-GFP cells exhibit similar morphology (E) Western blots of WM266-4-H2-GFP cell lysates.

### Development of primary HCS assay for hnRNPH2 down regulators

#### Choice of a readout

GFP fluorescence can be measured in HTS mode by either microplate readers or high content imagers. Microplate readers are preferable for HTS since they are ubiquitous and measurement requires less optimization, whereas high content imagers are more specialized, require fine tuning, and not always available at various HTS facilities. To determine whether an HTS assay can be developed using microplate reader we measured GFP fluorescence of range of WM266-4-H2GFP cells using Nunc® black opaque 384 well plate (Thermo Scientific cat# 264705). As can be seen in Fig. 2A, only 1,250 cells/well resulted in sub-confluent culture, which is a necessary condition for performing biological experiments. Ratio of GFP fluorescence intensity (FI) of GFP and non-GFP WM266-4 cells was 1.3 (Fig. 2B) suggesting that the assay window is very narrow and would not allow for separation of noise and hits. Since reader-based measurement approach did not produce desired signal separation, we proceeded with assay development using high content imager (CellInsight CX7, ThermoFisher Scientific). We utilized black clear-bottom 384 well plates (Greiner cat# 788091). These plates have the same well surface area as Nunc plates that we used for cell number determination (Fig. 2A), therefore, we used 1,250 cells/well of both WM266-4 and WM266-4-H2GFP cells for our experiments. High Content Assays (HCA) rely on algorithms that recognize and draw borders (masks) around objects that need to be quantitated. To enable a GFP fluorescence intensity measurement we developed a method based on a standard algorithm available as a part of CX7 software package. As can be seen in Fig. 2C, our method recognizes nuclei expressing hnRNPH2-GFP very well. This suggested that our algorithm can recognize valid objects and discarding non-specific signals.

**Figure 2.**
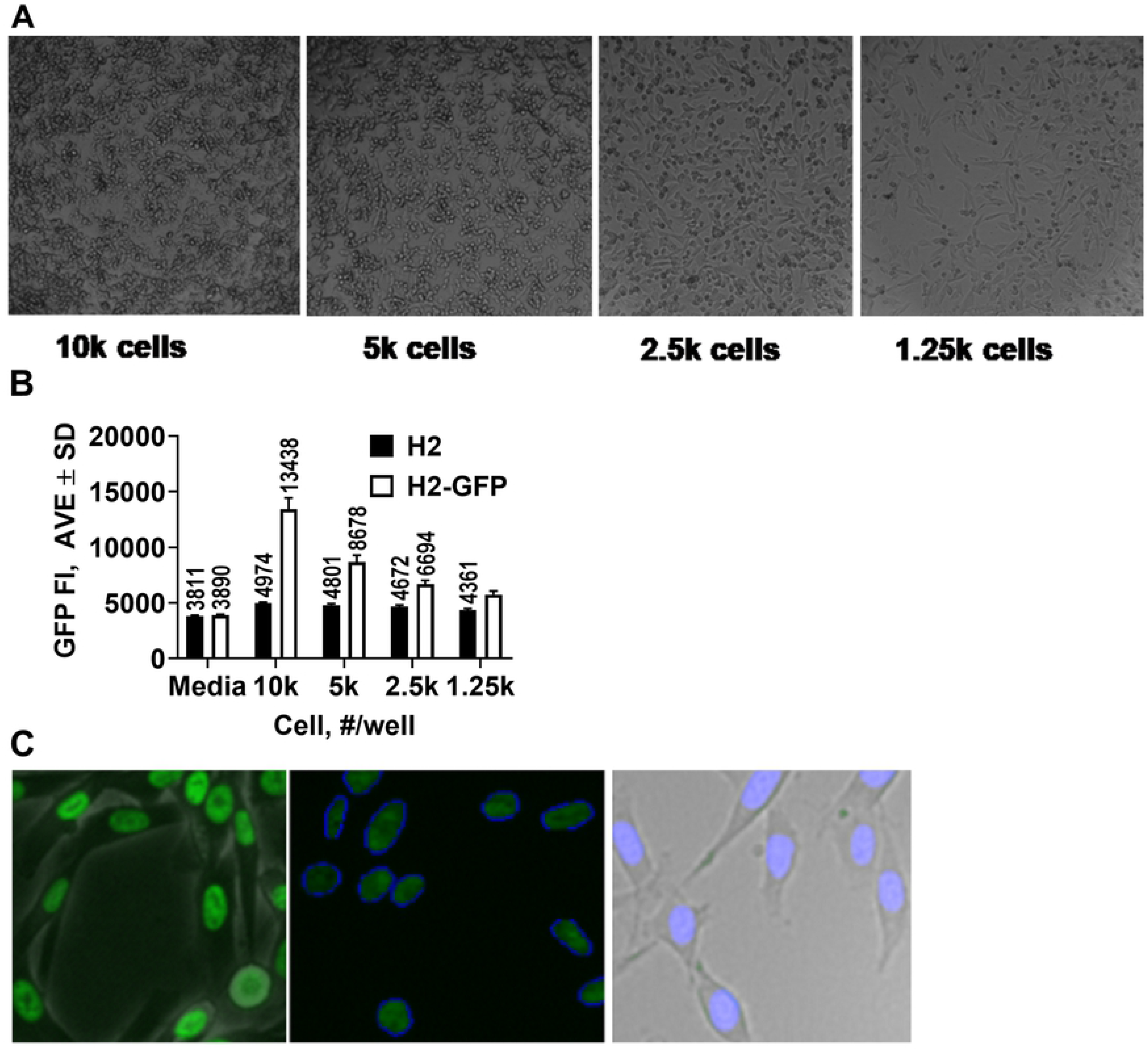
Selection of a readout approach. (1) WM266-4-H2-GFP cell number titration in 384 well plate black opaque plates. All panels imaged at 5x magnification; (B) Fluorescence intensity readout calculations of WM266-4-H2-GFP and WM266-4 cells in 384 well plate black opaque plates using a Biotek H1 plate reader; (C) WM266-4-H2-GFP cells imaged using ThermoFisher CellInsight CX7 instrument. Left panel – overlay of bright field and GFP channel; middle panel – GFP channel image of H2-GFP-expressing nuclei. Blue contour shows that the developed algorithm recognizes the shape of GFP-expressing nuclei for further quantitative analysis; right panel - overlay of bright field and DAPI channel. All panels imaged at 20x magnification.

Next, we dispensed 1,250 WM266-4-H2GFP cells well in 10μL of EMEM into each well of columns 2-24 of black clear-bottom 384 well plates (Greiner cat# 788091) and 1,250 WM266-4 cells in each well of column 1 and estimated signal-to-basal (S:B) ratio. As expected, wells with WM266-4 cells did not produce a signal while wells with WM266-4-H2GFP cells had Average Total Fluorescence Intensity (ATOI) of 338,408±15382 (n = 368 wells) (Fig. 3A). %CV was 4.5 suggesting great intra-plate signal reproducibility. While we could not calculate a true S:B ratio due to the lack of low ATOI control, the high signal of wells with WM266-4-H2GFP cells (high ATOI control) and a robust %CV suggested that a high content imager can be used to develop an HTS assay.

**Figure 3.**
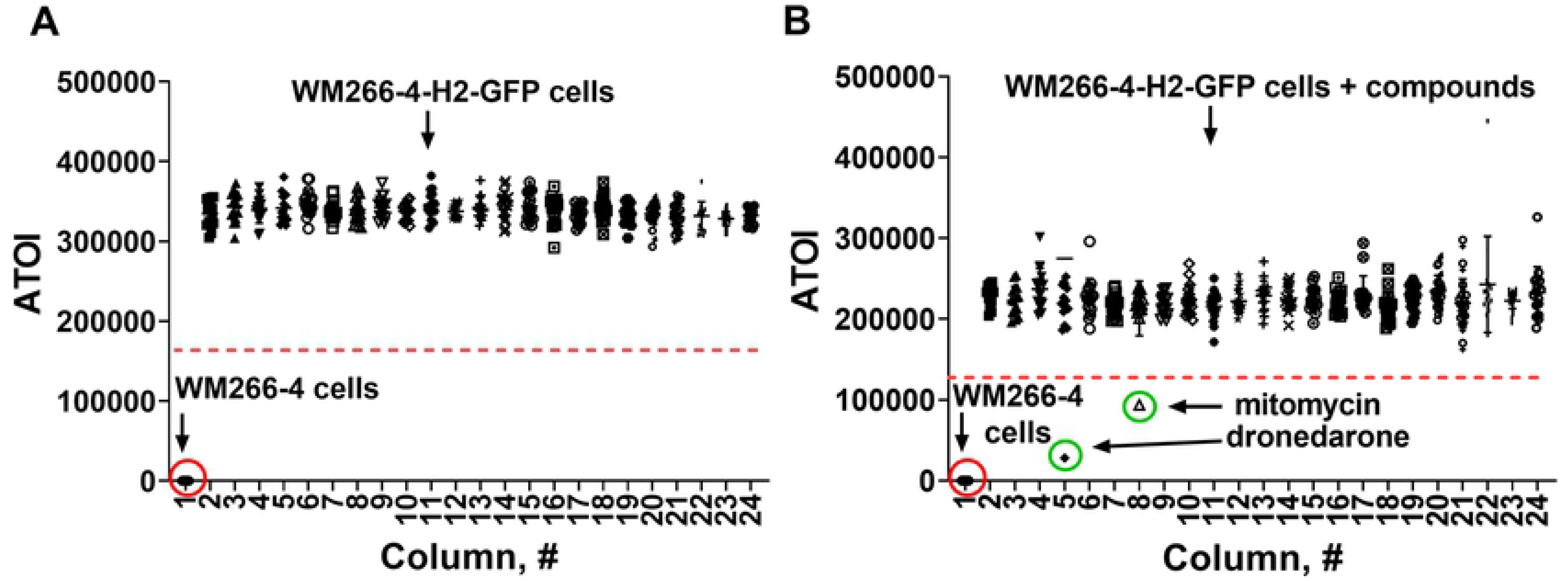
HCS assay 384 well plate scatter plots with and without test compounds. (A) Plate without compounds shows low scatter and large signal window between WM266-4-H2-GFP and WM266-4 cells. (B) Plate with test compounds shows 2 hits (green circle) that decreased GFP signal by more than 50%. Red punctured line indicates hit cutoff (50% signal reduction).

#### Identification of a pharmacological control for HCS assay

We conducted a screen of FDA-approved drug library (1,400 compounds) to identify a pharmacological control for HCS assay. Assay protocol followed “addition only” format (no extraction, filtration, or washing steps): (1) addition of 1,250 cells in 10 μL of EMEM media with 10% FBS, 1% PenStrep, 50 ug/mL hygromycin, utilizing Integra ViaFlo 384 liquid dispenser, (2) addition of test compounds (10μM final) using pin tool device (VP Scientific), (3) incubation of cells and compounds for 24 hrs at 37 °C, 5% CO_2_ and 95% relative humidity., and (4) measurement of Average Total Fluorescence Intensity using CellInsight CX7 (ThermoFisher) high content imager. We used 50% signal decrease as a hit cutoff (Fig. 3B).

Using this protocol, we found 8 hits (0.5% raw hit rate) of which we were able to purchase six compounds. Out of six commercially available compounds, four confirmed activity in dose response format (0.66% confirmed hit rate), namely dronedarone, mitomycin C, lapatinib free base form, and sunitinib (Fig. 4A-D). Two compounds exhibited EC_50_ values <20μM (0.07% high activity hit rate) (dronedarone = 17μM and mitomycin = 7.3μM).

**Figure 4.**
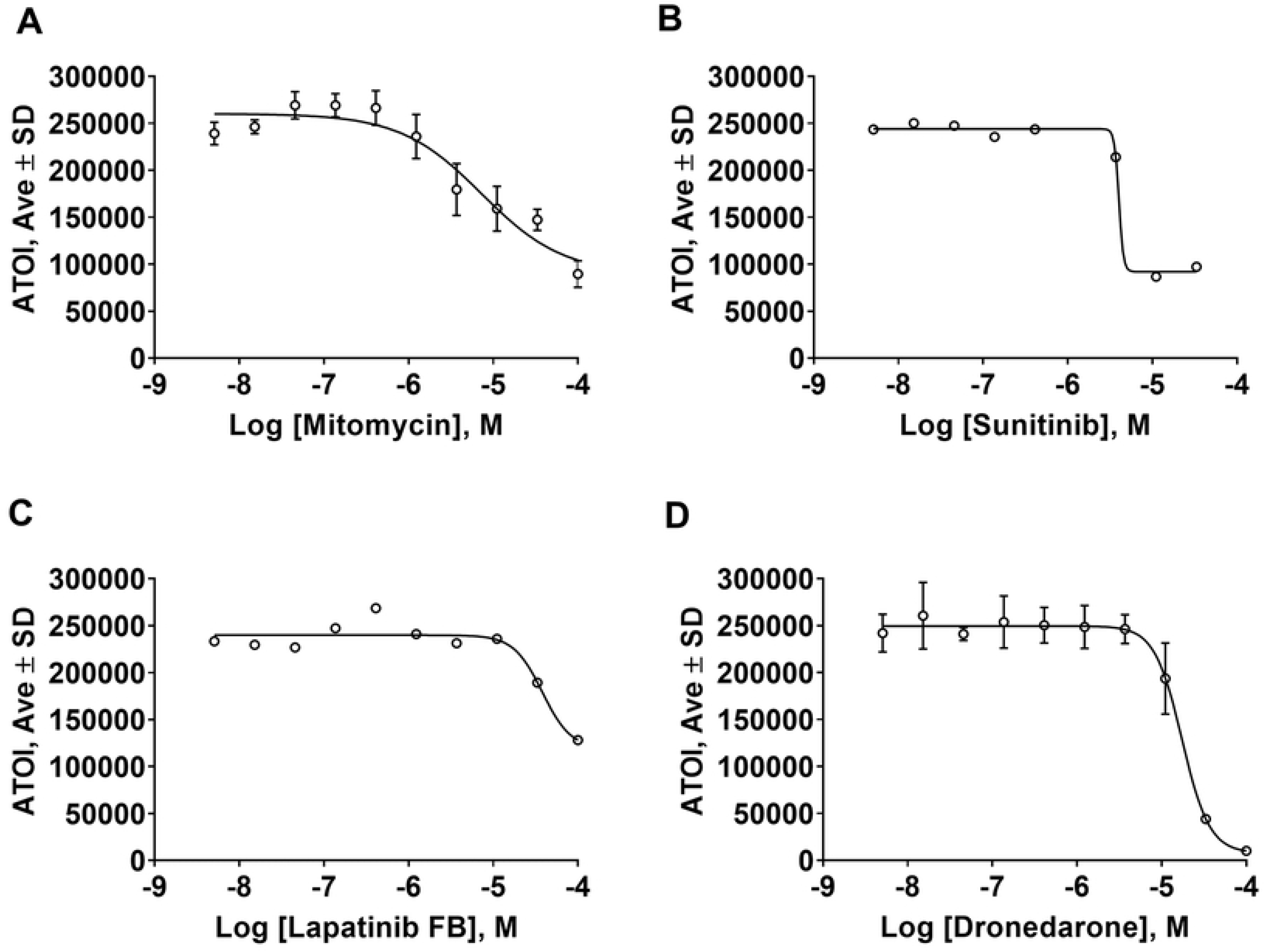
Dose response studies with pilot screen hits. Dose response studies in HCS assay for hits that show a pharmacological response. Lapatinib FB – lapatinib free base.

Examination of images of WM266-4-H2-GFP cells in presence of mitomycin showed a dose dependent decrease of fluorescence and nuclei shapes suggesting that HCS assay is capable of correctly recognizing and quantitating the objects in the wells (Fig. 5A). Interestingly, none of FDA approved melanoma drugs (Fig. 5B, trametinib, dabrafenib, vemurafenib, dacarbazine) decreased hnRNPH2 signal further corroborating our hypothesis that hnRNPH2 controls pathways previously unexplored for melanoma therapy (3).

**Figure 5.**
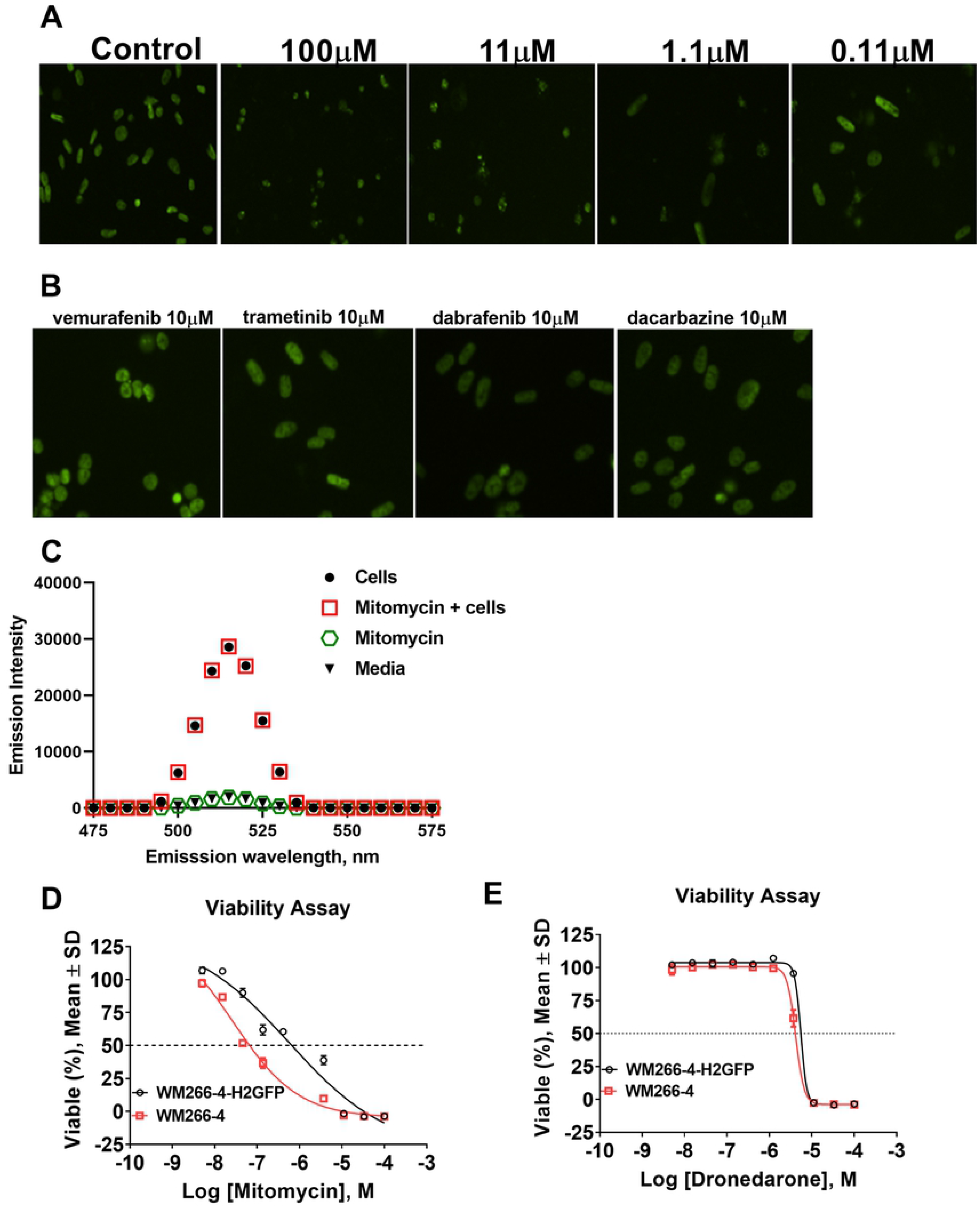
Dose response studies with dronedarone and mitomycin. (A) Images of WM266-4-H2GFP cells in the presence of varying concentrations of mitomycin show a dose-dependent change in GFP fluorescence. Images taken at 10x magnification using CellInsight CX7 instrument; (B) Images of WM266-4-H2GFP cells in the presence of FDA-approved melanoma drugs show no decrease of GFP fluorescence as compared to control (5A). Images taken at 10x magnification using CellInsight CX7 instrument. (C) Fluorescence emission scan of mitomycin in presence of WM266-4-H2GFP cells show no effect on fluorescence of GFP; Dose response studies in CellTiter Glo viability assay for (D) mitomycin and (E) dronedarone show that both compounds decrease viability of WM266-4 and WM266-4-H2GFP cells in a dose-dependent manner.

Compounds that absorb light in the GFP emission range (~500nm) could potentially be detected as hits in our assay format, therefore, we examined mitomycin for interference with GFP signal. Fluorescence emission scans of 10μM mitomycin in the presence and absence of WM266-4-H2-GFP cells showed no effect on GFP fluorescence (Fig. 5C) suggesting that mitomycin is a non-artifactual hit.

Next, we evaluated the effect of hits on melanoma cell viability. Interestingly, mitomycin was ~20-fold more potent against WM266-4 cells than WM266-4-H2-GFP cells in viability assay (Fig. 5D, IC_50_ = 27nM *vs* 570nM, respectively), whereas dronedarone was equipotent (Fig. 5F, IC_50_= 5μM). This result suggests different mechanism of action of mitomycin and dronedarone.

#### Assay validation in 384 wpf

Using 100μM dronedarone as a control we performed a small screen (n=3 plates) to validate assay robustness. All plate-based QC parameters were acceptable: %CV = 6.7±0.3, S/B = 21±2.1; Z = 0.5±0.02, Z’ = 0.75±0.04. Additionally, EC_50_ values of mitomycin and dronedarone were reproducible plate-to-plate (6.5±0.4 and 17±1.5μM, respectively).

#### Validation of hits using western blot

To confirm that signal decrease in HCS assay is authentic (i.e. due to the actual decrease of hnRNPH2-GFP expression and not due to the artifact such as spectral activity), we tested for the effect of 100μM mitomycin and dronedarone on hnRNPH2 levels in both transfected and parental cells using western blot. As evidenced by Fig. 6, mitomycin treatment partially decreased levels of hnRNPH2-GFP, hnRNPH2 and H1 in both cell lines (Fig.6-C), whereas dronedarone treatment resulted in a complete loss of hnRNPH2 and H1 in WM266-4 cells (Fig. 6A and D). These results correlate well with the dose response in HCS assay (Fig. 4AB) where mitomycin showed partial pharmacological response and dronedarone showed full response.

**Figure 6.**
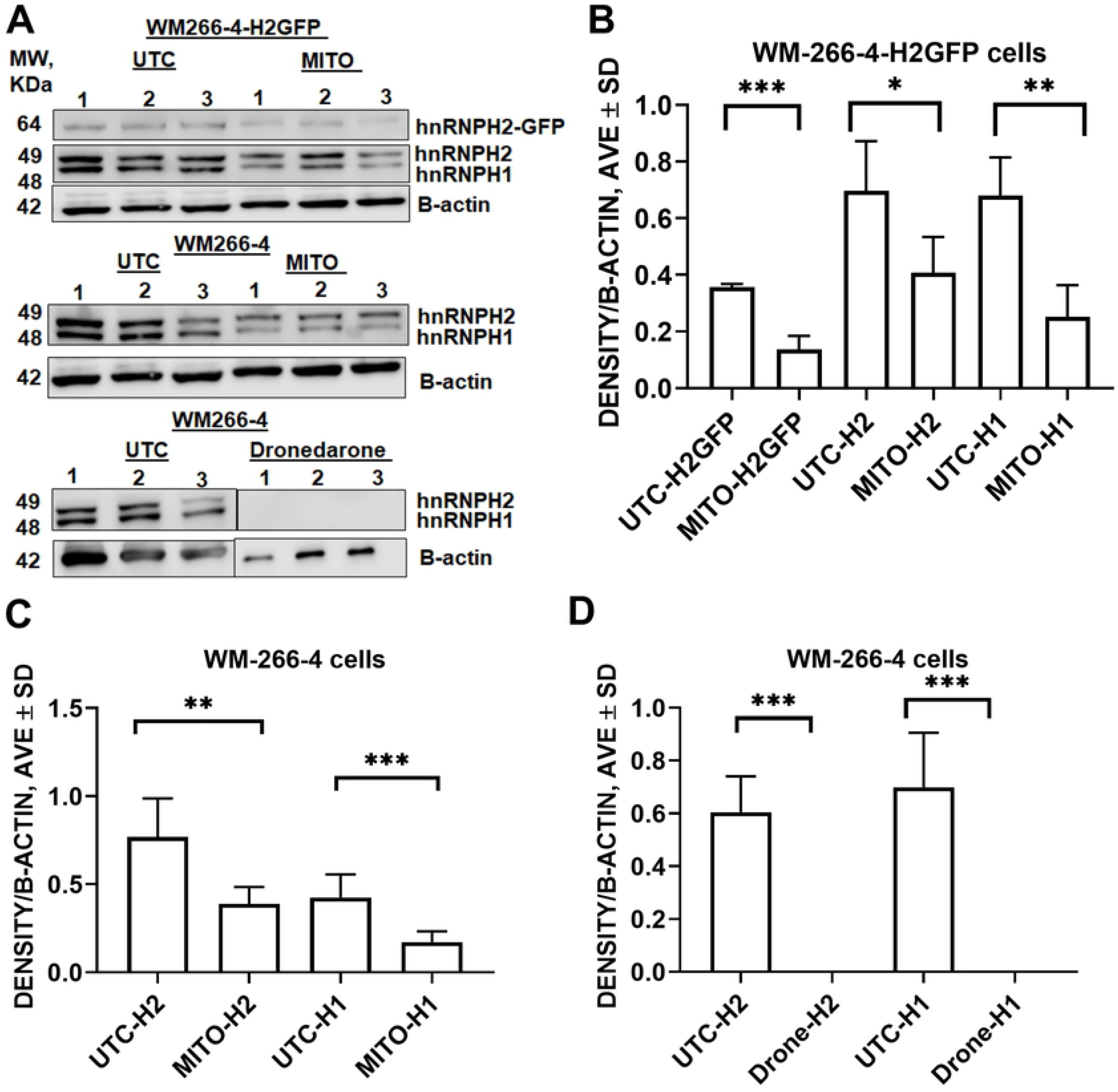
Authentication of pilot screen hits dronedarone and mitomycin using western blot. (A) Western blots and (B-D) western blot quantitation of WM266-4 and WM266-4-H2GFP cell lysates after 24 hr incubation with 100μM mitomycin and dronedarone. The data shown are the mean ± SD, n=3. ***** - p < 0.0001, *** - p < 0.001, ** - p < 0.01, * - p < 0.05. Mito – mitomycin, H1 – hnRNPH1, H2 – hnRNPH2, UTC – untreated control, drone – dronedarone.

#### Assay miniaturization to 1,536 well plate

To enable future HTS campaigns, we scaled down the assay final volume in 1,536 well plate format by factor of 2. Since fewer number cells is used in 1,536 wpf than 384 wpf (750 vs 1,250, respectively) we ascertained that we can detect sufficient number of cells per field of vision (Fig. 7A). We observed that normalization of Average Total Object Intensity (ATOI) by the cell number produces lower scatter, therefore, we used ATOI/cell number ratio as a basal result for calculations. Next, we conducted mitomycin and dronedarone dose response study in 1,536 wpf. EC_50_ values of mitomycin and dronedarone were reproducible between 384 and 1,536 wpf (Fig. 7B and C; 17μM and 15μM for dronedarone in 384 and 1,536 wpf, respectively; 6.5μM and 15μM for mitomycin in 384 and 1,536 wpf, respectively). All plate-based QC parameters were acceptable: %CV = 14, S/B = 84; Z = 0.5, Z’ = 0.52 suggesting readiness of the assay to the screening.

**Figure 7.**
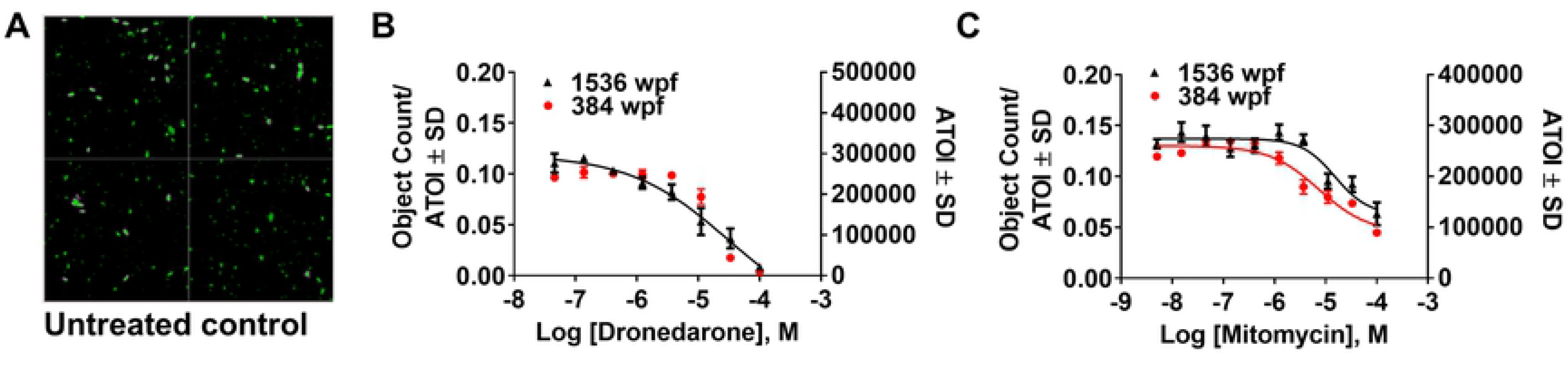
Assay miniaturization to 1,536 well plate format. (A) Image of WM266-4-H2GFP cells in 1,536 well plate. Image taken on GFP channel using 10x magnification. Four fields of vision were acquired; Comparison betweeen 384 wpf and 1,536 wpf using (B) dronedarone and (C) mitomycin dose response study. 384 wpf data plotted on the right Y-axis, 1,536 wpf data plotted on the left Y-axis.

Next, we performed the assay in triplicate to validate its robustness. All plate-based QC parameters were acceptable: %CV = 6.7±0.3, S/B = 41±11; Z’ = 0.54±0.09 suggesting readiness of the assay to the large-scale screening.

## DISCUSSION

Our previous studies show that decrease of hnNRPH2 and H1 expression as a result of binding by a small molecule leads to the selective death of melanoma cells (3). This suggested that hnRNPH2 and associated pathways can be targeted for melanoma therapy; therefore, we decided to expand small molecule compound tools available *via* HTS effort in order to explore such pathways. We reasoned that compounds that decrease of hnNRPH2 and H1 expression do not necessarily have to bind it. Therefore, the primary biochemical assay that detects just hnNRPH2 binders would be unnecessarily excluding compounds that decrease hnNRPH2 expression *via* hitherto unknown mechanisms, but could be just as valuable. Based on these considerations we developed more physiologically relevant assay that can easily monitor level of hnNRPH2 protein inside the cells in the presence of test compounds.

Most of HTS-amenable detection technologies are based on either luminescence or fluorescence measurements, however, HTS libraries contain ~3% luciferase inhibitors (13, 14) and spectrally active compounds (15). Since we are interested in compounds that decrease assay signal, luciferase inhibitors would result in ~3% false positives. On other hand, fluorescent compounds could mask some true hits by artificially increasing fluorescence (false negatives) but would not produce false positives. Quenchers of fluorescence are much rarer, therefore, there is less concern about false positives when using fluorescence rather than luciferase reporter. For example, there estimated to be ~0.01-0.2% of compounds with Ex_480nm_/Em_535nm_ (green fluorescence) (14). Indeed, none of the pilot screen hits affected GFP fluorescence suggesting that our assay is not prone to fluorescent artifacts possibly due to the high quantum yield of SparkGFP.

The purpose of developing of an assay for hnRNPH2 is to discover novel down regulators of its expression working via different mechanisms. This, in turn, will help researchers to gain an insight into pathways and mechanisms that depend on hnRNPH2 expression. Two confirmed hit compounds discovered as a result of screen of FDA library, dronedarone and mitomycin C, have not been described previously in connection with hnRNPH2 and, therefore, could be valuable chemical tools to study its biology. Mitomycin C is a pro-drug that can be activated intracellularly by several oxido-reductases whose expression is often elevated in cancers (16). Multiple mechanisms by which mitomycin C kills cancer cells have been proposed, including formation of DNA and RNA adducts (17). Global expression profiling of U2OSE64b cells treated with mitomycin C showed that it affected mRNA splicing and transport (18), which is consistent with what is known about hnRNPH2 role (19) in cellular processes. Mitomycin is used in clinic for various indications including conjunctival melanoma (20). Mitomycin was less potent against WM266-4-H2GFP cells that overexpress hnRNPH2 and hnRNPH1 (Fig. 5D), suggesting that its effect was dependent on hnRNPH2 and hnRNPH1 concentration. This potentially means that in addition to nucleic acid modification, mitomycin acts by binding hnRNPH2 directly, which was not previously described. This hypothesis is corroborated by the fact that 2155-14, a known hnRNPH2-binder (3), exhibits the same trend in inhibiting of WM266-4 and WM266-4-H2GFP cells (Fig. 1E).

Dronedarone is a multi-ion channel blocker used in the clinic as an anti-arrhythmic medicine (21). It was active against ovarian (22) and breast cancer cells *in vitro* and *in vivo* (23), however, its activity against melanoma was not previously demonstrated. However, another calcium channel blocker, amlodipine, inhibited uveal malignant melanoma and cutaneous malignant melanoma cell lines with IC_50_ values of 13.1 μM and 15.9 μM, respectively (24), which is similar to dronedarone’s potency (17±1.5μM) against cutaneous malignant melanoma (WM266-4). Based on dronedarone’s equipotent inhibition of WM266-4 and WM266-4-H2GFP cells (Fig. 5E) it is reasonable to hypothesize that its effect is not modulated by hnRNPH2 binding. It is also possible that hnRNPH2 down regulation is a consequence of ion channel blockade by dronedarone.

### Conclusions

As a result of the studies described herein, we developed novel High Content Assay for down regulators of hnNRPH2 amenable to the high throughput screening. A screen of FDA drug library identified two compounds that down regulate hnNRPH2 *via* dissimilar mechanisms and inhibit viability of malignant melanoma cells. Further studies are needed to dissect the mechanistic differences of hit compounds’ effects. In summary, our novel assay is suitable platform for large scale screening for novel down regulators of hnNRPH2. To our knowledge, this is a first HTS/HCS assay that allows monitoring of hnNRPH2 protein expression in live cells.

## ACKNOWLEDGMENT

Authors thank Drs. Brent Samson and Steven T. Frank (ThermoFisher) for help with CX7 method development.

## Supporting Information

### AUTHOR INFORMATION

Correspondence to: Dmitriy Minond, Rumbaugh-Goodwin Institute for Cancer Research, Nova Southeastern University, 3321 College Avenue, CCR r.605, Fort Lauderdale, FL 33314;

### AUTHOR CONTRIBUTIONS

DM envisioned and designed the study, performed HCS assay development and pilot screen experiments, analyzed all experimental data, and wrote the manuscript. JD developed WM266-4-H2GFP cell line, SB and PS performed hit validation western blot experiments. SR, TPS and LSD performed 1,536 well plate experiments. All authors have given approval to the final version of the manuscript.

### FUNDING SOURCES

This work was supported by the National Institutes of Health (AR066676 and CA249788 to DM).

